# Brain2VLM: Hierarchical Alignment Between Cortical Representations and Vision-Language Latent Spaces

**DOI:** 10.64898/2026.04.23.720313

**Authors:** N. A. Adarsh Pritam, Jeba Shiney O, Sanyam Jain

## Abstract

This work introduces Brain2VLM, a framework for analyzing how cortical representations align with latent spaces of pretrained diffusion-based vision-language models for brain-to-image reconstruction. While recent approaches achieve strong performance by mapping functional Magnetic Resonance Imaging (fMRI) signals to model latents, the structure of this mapping remains poorly understood. We hypothesize that brain-to-latent alignment is hierarchical, with early visual cortex exhibiting approximately linear correspondence to structural diffusion latents, and higher-order visual areas requiring nonlinear mappings to align with semantic embedding spaces. To test this, we decode diffusion latents and CLIP embeddings from fMRI signals across four subjects from the Natural Scenes Dataset (NSD) using both linear ridge regression and a nonlinear residual MLP. Our results reveal that nonlinear decoding provides only marginal improvements for diffusion latents (*Δ* ≈ 0.05 − 0.06 in correlation), but yields substantial gains for semantic embeddings (*Δ* ≈ 0.47), significantly improving distributional alignment (MMD: 0.042 vs 0.358). However, increased decoder expressivity can introduce shifts in latent distributions, highlighting a trade-off between prediction accuracy and generative compatibility. Despite using a simple reconstruction pipeline, Brain2VLM consistently improves semantic reconstruction quality across all four subjects while maintaining competitive pixel-level fidelity (PixCorr 0.36, CLIP 85%), suggesting that improvements in brain-to-latent alignment play an important role in reconstruction quality alongside generative modeling. These findings are consistent with a hierarchical pattern of alignment between cortical representations and model latent spaces, highlighting the brain-to-latent interface as an important component of brain decoding systems. Our code can be found at https://github.com/adarsh-crafts/Brain2VLM

## 1 Introduction

Understanding how the human brain represents and reconstructs visual experience remains a central challenge in NeuroAI. Brain decoding aims to recover sensory stimuli from non-invasive brain recordings such as fMRI, enabling the reconstruction of perceptual and semantic content from neural activity. Earlier studies trained new generative models with fMRI data from scratch, or fine-tuned toward the specific stimuli used in the fMRI experiments [11,19,21]. While they showed impressive results, they achieved limited success in pixel-wise and semantic fidelity. This limitation arises partly from the small number of samples available in neuroscience datasets and from the difficulties involved in the training of complex generative models [20]. Diffusion models, particularly latent diffusion models (LDMs), achieve state-of-the-art performance in conditional image generation by operating in a compressed latent space and conditioning the denoising process on semantic embeddings [6, 17]. These representations capture perceptual and semantic structure, making them suitable for brain decoding, where neural activity is more likely to align with abstract representations rather than raw pixel intensities. Recent work, illustrated in Fig. 1 (middle row), shows that high-resolution (512 × 512) images can be reconstructed from brain activity using simple linear decoders without training or fine-tuning the generative model [20].

**Fig. 1:**
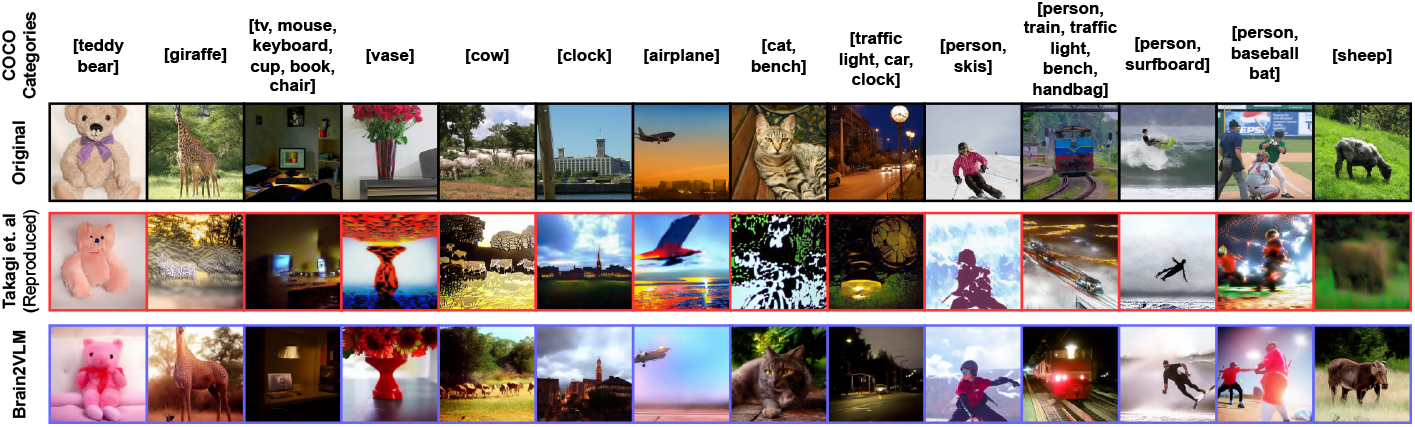
Columns show stimuli; rows show (black box, top row) presented images (drawn from the publicly available MS COCO dataset [12]), (red box, middle row) images reconstructed from fMRI signals using linear decoders, and (purple box, bottom row) reconstructions using proposed nonlinear decoders for representative reconstructions from subj01. While both methods recover coarse spatial structure, nonlinear decoding improves semantic consistency, yielding more recognizable object categories (e.g., animals, vehicles). However, these semantic improvements do not consistently translate into sharper low-level details, highlighting a gap between semantic alignment and pixel-level reconstruction quality.

In such approaches, neural activity patterns are first mapped to latent representations of pretrained generative models using relatively simple decoders such as linear regression or neural networks. These decoded latent representations are then used as conditioning signals for the diffusion model, which reconstructs the image through an iterative denoising process. This separation allows leveraging strong generative priors while simplifying the decoding task.

Although recent models achieve increasingly impressive reconstruction accuracy, performance gains alone do not clarify a deep question: what information is actually represented in the cortical activity during vision? Does brain activity encode detailed pixel-level information, or does it reflect abstract, latent semantic and perceptual structure? A growing field of neuroscience research suggests that the brain does not store literal pixel-level snapshots of scenes [10, 15]. Instead, distributed patterns of activity across visual and higher-order cortex are systematically related to high-level and semantic features, including object categories and conceptual structure, rather than the exact low-level sensory details [7, 9].

In agreement with this perspective, recent brain decoding studies increasingly rely on mappings from fMRI signals into high-level latent spaces of deep neural networks, which organize visual information according to semantic similarity [11, 13, 16–18]. Interestingly, as shown in Fig. 1 (middle row), even simple linear mappings preserve semantic content despite deviations in low-level detail. Nonlinear decoding further improves the recognizability of high-level semantic attributes, while low-level properties such as texture remain largely unchanged, suggesting that improved latent alignment does not necessarily translate into perceptual gains.

However, it remains unclear whether cortical representations align uniformly with the latent spaces of modern generative models, or whether this relationship depends on the level of abstraction in both brain and model representations. We hypothesize that brain-to-latent alignment is hierarchical: early visual areas exhibit approximately linear mappings to structural diffusion latents, while higher-order visual areas require nonlinear mappings to align with semantic embedding spaces such as CLIP. Using the ROI partition established by [20], we investigate this hypothesis by decoding diffusion latents and CLIP embeddings from four NSD participants using both linear ridge regression and a nonlinear residual MLP. We then assess whether the observed alignment patterns generalize beyond a single individual.

Unlike much of the prior work that primarily improves reconstruction through complex generative pipelines and conditioning mechanisms [3, 4, 13, 16, 18], we isolate and study the role of the brain-to-latent mapping itself, which allowed us to investigate whether its structure depends on representational hierarchy. Our contributions are:

1. A hierarchical analysis of brain-to-latent alignment showing that early visual cortex exhibits approximately linear mappings to structural diffusion latents, while higher-order visual areas require nonlinear mappings to semantic embedding spaces.
2. Empirical evidence across multiple NSD subjects that nonlinear decoding yields substantial improvements for semantic representations (CLIP embeddings) but only marginal gains for structural diffusion latents, revealing distinct alignment regimes.
3. An analysis of the trade-off between latent prediction accuracy and reconstruction quality, showing that increased decoder expressivity can distort latent distributions despite improving correlation.
4. A simple residual MLP decoder that exposes these effects and provides a comparatively stronger brain-to-latent interface compared to standard linear regression.

## 2 Related Works

Early works in visual decoding focused on stimulus identification and constrained reconstruction [8, 14]. Object and action categories are organized within a continuous semantic space distributed across cortex [7]. [11] projected fMRI signals into CLIP-aligned embeddings to reconstruct complex natural scenes using GAN-based generation. [16] proposed a two-stage latent diffusion framework combining very-deep variational auto-encoder (VDVAE) structure prediction with diffusion-based refinement, and [20] demonstrated that high-resolution reconstructions could be obtained by linearly mapping fMRI signals into Stable Diffusion (SD) latent components without training or fine-tuning the generative model. Subsequent work extended this paradigm using multimodal decoding and CLIP-aligned representations, including caption-and-image reconstruction [3, 4], diffusion-prior approaches [18], and CLIP-based neural alignment [13]. These results suggest that brain decoding is most effective when mapping neural activity into structured latent spaces of pretrained generative models.

## 3 Methodology

### 3.1 Dataset

We used the Natural Scenes Dataset (NSD) [1], a large-scale 7*T* fMRI dataset in which participants viewed natural scene images during repeated scanning sessions. The stimuli consist of photographs drawn from the Microsoft COCO dataset [12]. Following prior brain decoding studies [16,20], we evaluated the proposed framework on four NSD participants (subj01, subj02, subj05, and subj07), which completed the full experimental protocol. Each subject viewed approximately 10,000 unique images across 30–40 sessions, with multiple repetitions for a subset of stimuli. Following prior work, we evaluated the proposed framework on four NSD participants (subj01, subj02, subj05, and subj07), all of whom completed the full experimental protocol. Each participant viewed approximately 10, 000 unique images across 30–40 sessions. Models were trained and evaluated independently using the standardized NSD train/test split, consisting a total of 982 shared test images across subjects and the remaining trials were used for training. For the test set, responses across repetitions were averaged, while individual trials were used for training. For functional data, we used the preprocessed single-trial beta weights provided by the NSD dataset, estimated using a denoising GLM with fitted hemodynamic response functions, as described in [1].

Voxel responses were extracted from visual cortex regions of interest including early visual areas (*V* 1–*V* 3) and higher visual areas (*V* 4, LO, PHC). We adopt the ROI partition introduced by [20]: early visual areas (V1–V3) are used for diffusion latent prediction, while higher visual areas (V4, LO, PHC) are used for CLIP embedding prediction. This separation allows testing whether alignment depends on representational level. To provide an overview of the Brain2VLM pipeline, we illustrate the full reconstruction process in Fig. 2.

**Fig. 2:**
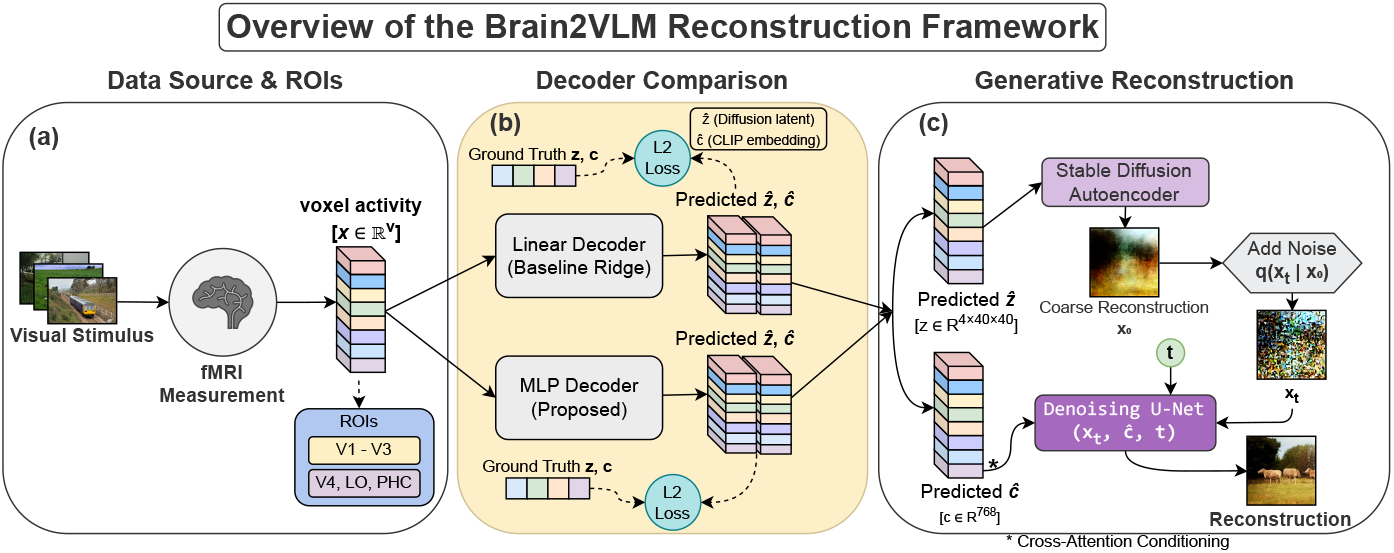
Brain2VLM reconstruction framework. (a) Visual stimuli are presented while fMRI responses are recorded from regions of interest (ROIs) in the visual cortex. Voxel activity from early visual areas (*V* 1–*V* 3) and higher visual areas (*V* 4, LO, PHC) forms the neural input representation. (b) Neural decoding models map voxel activity to latent representations of a pretrained diffusion model. We compare a linear Ridge decoder with our proposed nonlinear residual MLP decoder, both trained with *ℓ*_2_ loss to predict the diffusion latent 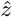 and CLIP embedding *ĉ*. (c) The predicted diffusion latent 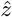 is decoded by the Stable Diffusion autoencoder to produce a coarse image *x*_0_. Noise is added via the forward diffusion process *q*(*x*_*t*_ | *x*_0_) and the image is refined through a denoising U-Net conditioned on *ĉ* via cross-attention, producing the final reconstruction.

### 3.2 Latent Diffusion Models

Image reconstruction was performed using Stable Diffusion v1.4, a latent diffusion model that generates images through iterative denoising in a compressed latent space [17]. Given a stimulus image *I* resized to 512 × 512, the pretrained model produces two latent representations as shown in equations 1 and 2.

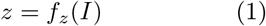

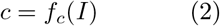

where *z* ∈ ℝ^4*×*40*×*40^ is the latent representation of the Stable Diffusion autoencoder and *c* ∈ ℝ^768^ is the CLIP image embedding used for cross-attention conditioning.

During reconstruction, the predicted diffusion latent 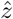 is first decoded by the Stable Diffusion autoencoder to produce a coarse image *x*_0_. Noise is then added through the forward diffusion process *q*(*x*_*t*_ *x*_0_) and the image is iteratively refined using a denoising U-Net conditioned on the predicted CLIP embedding *ĉ* via cross-attention. All parameters of the generative model remain frozen during training.

### 3.3 Decoding and Reconstruction Pipeline

Our goal is to reconstruct visual stimuli from fMRI signals by decoding neural activity into the latent representations of a pretrained diffusion model.

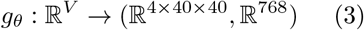

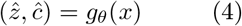

Let *x* ∈ ℝ^*V*^ denote the fMRI activity vector measured across *V* voxels for a single stimulus presentation. The decoding objective is to learn a mapping defined in Eq. 3 such that the predicted latent variables in Eq. 4 approximate the latent representations (*z, c*) extracted from the stimulus image.

We compare two neural decoding models that map voxel activity to diffusion latent representations: (1) a linear ridge regression decoder reproducing [20], and (2) a nonlinear residual multilayer perceptron (MLP) decoder. All decoder architectures and hyperparameters were kept identical across subjects.

1. **Linear decoder:** The baseline model learns a linear mapping between voxel activity *x* ∈ ℝ^*V*^ and latent representations *ŷ*, where *W* ∈ ℝ^*d×V*^ and *b* ∈ ℝ^*d*^. Parameters are estimated using ridge regression with an *ℓ*_2_ regularization penalty on *W* .
2. **Residual MLP decoder:** To capture nonlinear relationships between cortical activity and latent representations, we introduce a deep residual MLP decoder (Fig. 3).

**Fig. 3:**
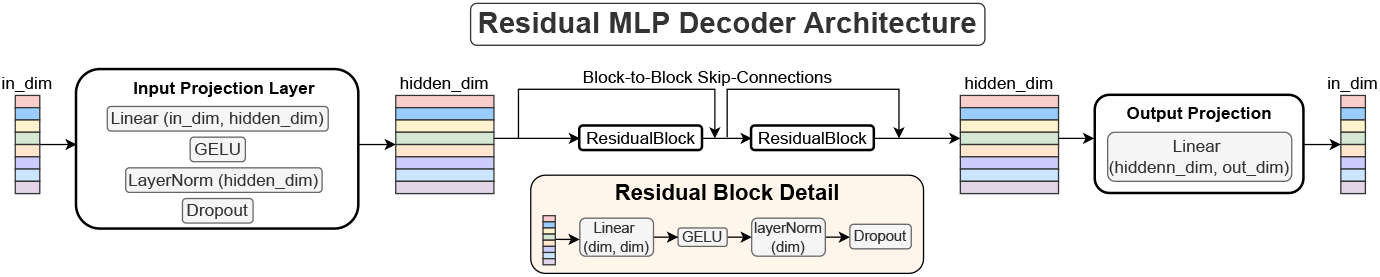
Residual MLP decoder architecture. The proposed decoder maps voxel activity to diffusion latent representations. An input projection first maps the voxel vector to a hidden representation using a linear layer followed by GELU, LayerNorm, and dropout. The hidden representation is processed through a stack of *N* residual blocks, each consisting of a linear layer, GELU activation, LayerNorm, and dropout with residual connections. A final linear layer projects the hidden representation to the target latent dimension. For illustration, the diagram shows *N* = 2 residual blocks, while the depth *N* is varied in our ablation experiments.

The voxel activity is first projected into a 2048-dimensional hidden representation, followed by *N* residual blocks. Each block consists of a linear transformation with GELU activation, LayerNorm, dropout, and a residual skip connection.

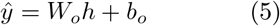

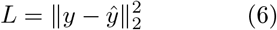

Finally, the hidden representation is projected to the latent space to produce the predicted latent vector *ŷ* (Eq. 5). Both the linear baseline and the MLP decoder are trained using the mean squared error loss defined in Eq. 6.

## 4 Results & Discussion

We evaluate the proposed decoder across four NSD subjects to determine whether improvements in brain-to-latent alignment generalize across individuals. We further analyze how these improvements impact reconstructed images, both qualitatively (Fig. 1) and quantitatively (Table 1).

**Table 1.**
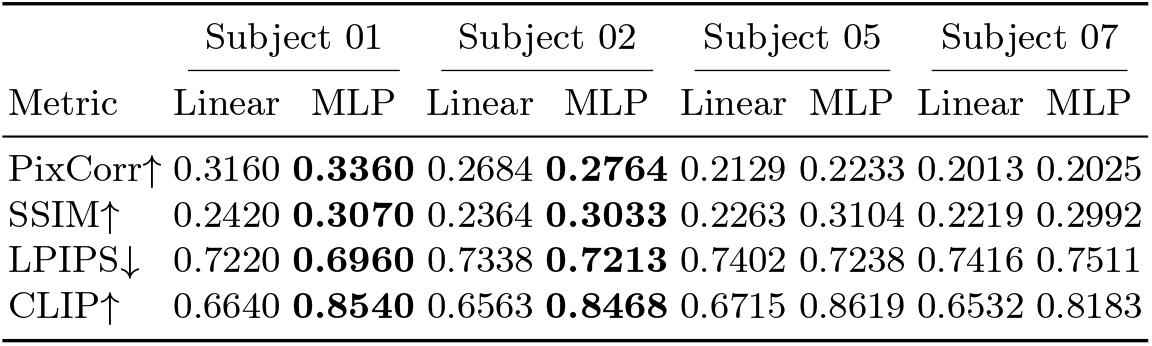
Effect of decoder choice on reconstruction quality using pixel correlation (PixCorr), Structural Similarity Index (SSIM), Learned Perceptual Image Patch Similarity (LPIPS), and CLIP-based semantic similarity.

### 4.1 Latent Prediction Accuracy

Figure 4 summarizes ablations over model depth, hidden width, and training data fraction. The decoder ablations are illustrated using subj01, while reconstruction experiments are reported across all four subjects. Increasing depth improves performance initially but saturates beyond two residual blocks (Fig. 4(i)), while width scaling yields consistent gains up to 2048 hidden units before plateauing (Fig. 4(ii)). Performance improves with additional training data (Fig. 4(iii)), indicating that decoding remains partly data-limited. Table 4b reports the best configurations, which are statistically significant (*p <* 0.01, paired t-test across stimuli).

**Fig. 4:**
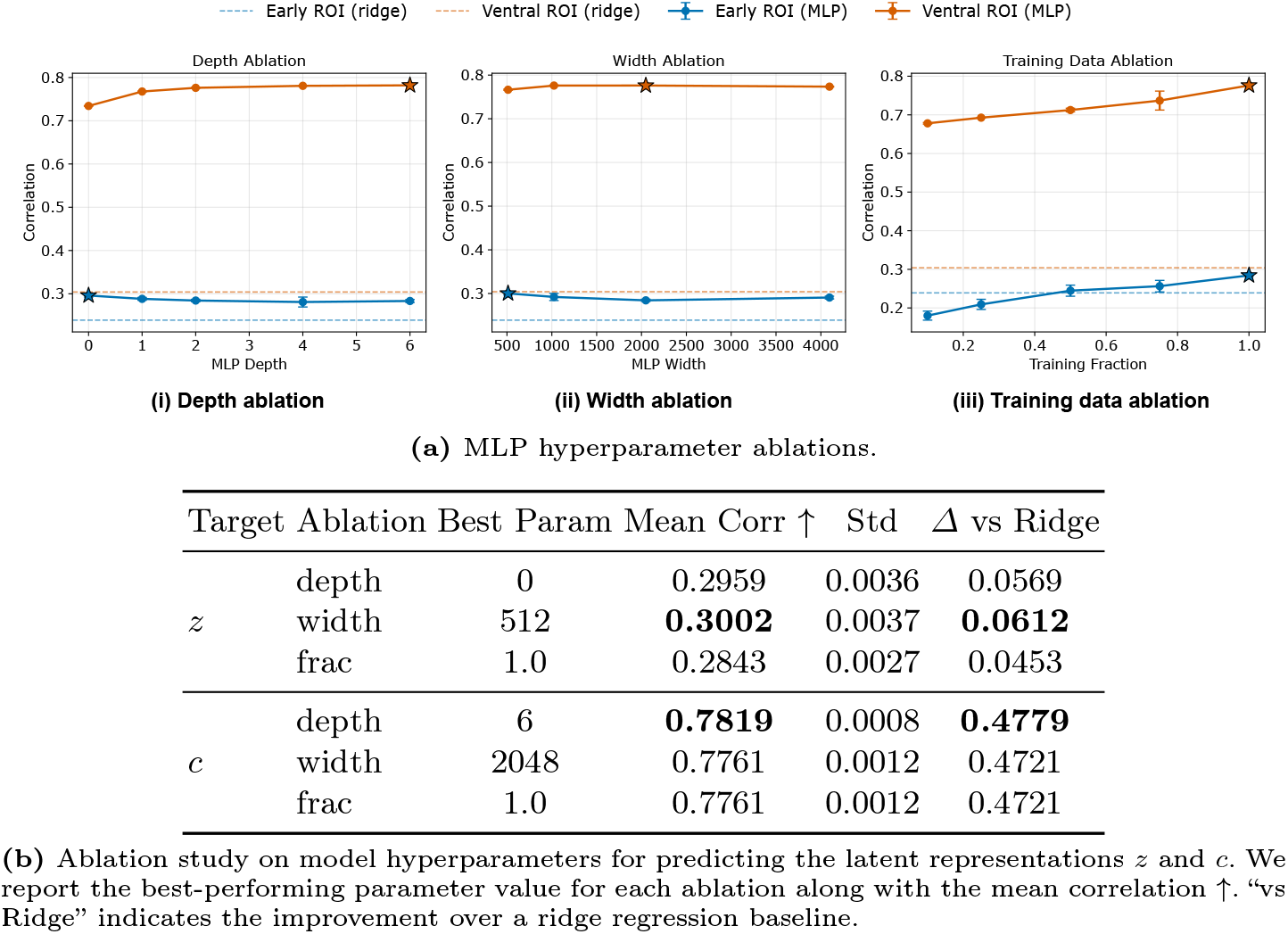
Residual MLP decoder ablation analysis. (a) Top: Effect of decoder depth, hidden width, and training data fraction on latent prediction accuracy. (b) Bottom: Quantitative summary of the best-performing configurations compared with a ridge regression baseline.

A clear divergence between structural and semantic latent targets is observed. For diffusion latents (*z*), nonlinear decoding provides only modest improvements over linear regression (*Δ* ≈ 0.05–0.06), suggesting that the mapping from early visual cortex to structural latent representations is approximately linear with some nonlinear corrections. In contrast, for semantic embeddings (CLIP, *c*), non-linear decoding yields substantial gains (*Δ* ≈ 0.47), indicating that mappings from higher-order visual areas to semantic latent spaces are strongly nonlinear.

Interestingly, we observe a systematic mismatch between latent prediction accuracy and reconstruction quality. This effect is also reflected qualitatively in Fig. 1, where nonlinear decoding improves the qualitative recognizability of high-level semantic content without consistently enhancing low-level visual detail. We hypothesize that this is due to distribution shifts introduced by the nonlinear decoder, which improve regression accuracy but reduces the compatibility with the generative prior. This suggests a fundamental limitation in current brain-to-image pipelines: accurate latent prediction is necessary but not sufficient for high-quality image reconstruction.

### 4.2 Reconstruction Quality

As shown in Table 1, ccross all four subjects, nonlinear decoding consistently improves semantic metrics while generally improving low-level reconstruction quality, with larger gains observed in semantic alignment than in pixel-level fidelity. The trend is consistent with the qualitative examples in Figure 1, where improvements are more apparent in semantic consistency (e.g., object identity) than in fine-grained visual structure. This suggests that improvements in latent prediction accuracy are reflected more strongly in semantic consistency than in fine-grained visual detail. Although the magnitude of improvement varies across participants, the overall trend remains consistent, indicating that the proposed nonlinear decoder generalizes beyond a single subject.

Figure 5 presents representative good, average, and challenging reconstruction examples. (1) Good cases: both decoders recover global structure; the MLP produces more semantically consistent object representations, indicating improved alignment with high-level latent structure. (2) Average cases: differences become more pronounced — linear reconstructions exhibit blurred or fragmented content, while the MLP preserves object identity and scene layout. (3) Challenging cases: linear decoding often collapses to noise or spurious textures, whereas the MLP retains coarse semantic organization despite loss of fine detail. These observations support the finding that improving brain-to-latent alignment primarily enhances semantic consistency rather than low-level fidelity. Notably, even when pixel-level quality is limited, better latent alignment yields more recognizable object categories, consistent with the large gains observed for CLIP embeddings.

**Fig. 5:**
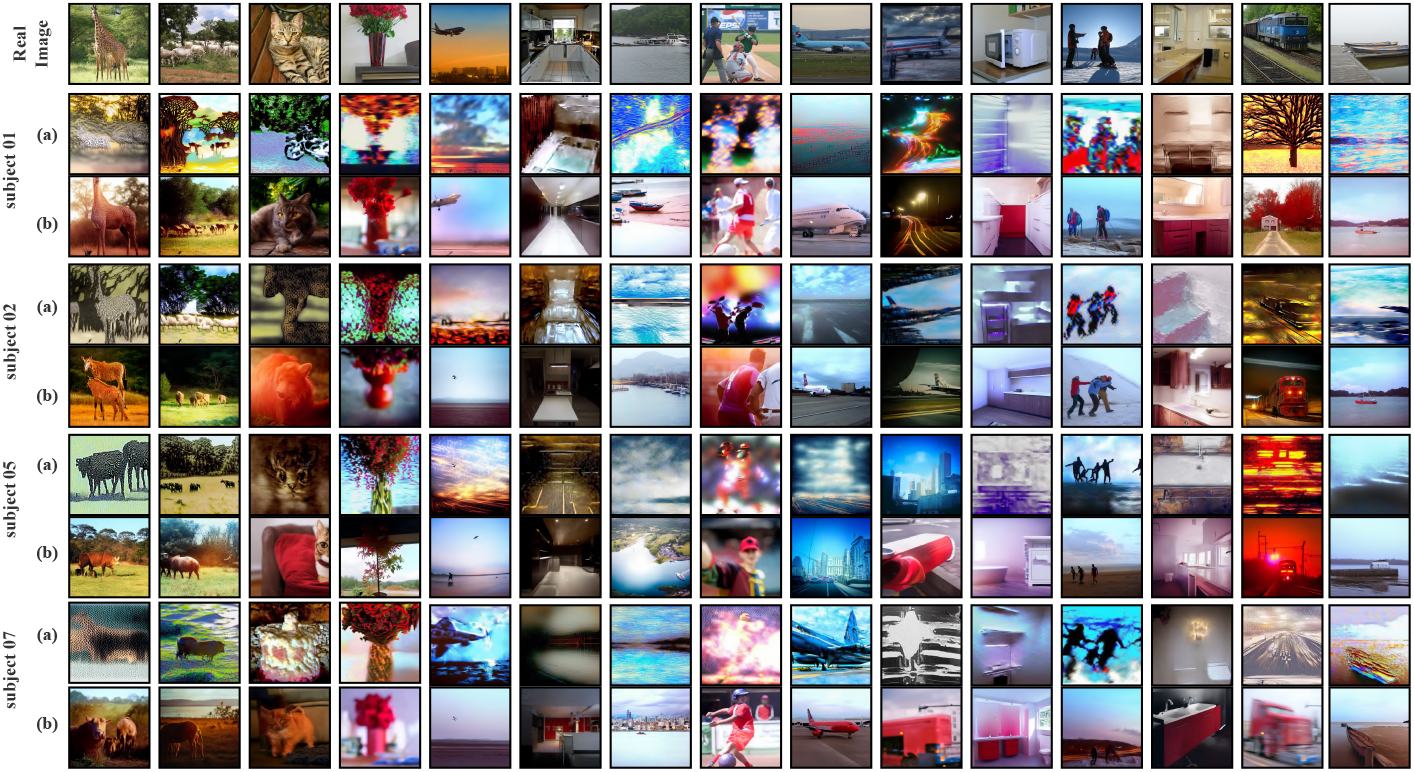
Qualitative comparison of image reconstructions across subjects. Columns correspond to different visual stimuli, while the top row shows the corresponding ground-truth images. For each subject (subj01, subj02, subj05, and subj07), row (a) presents reconstructions obtained using the linear ridge decoder, and row (b) shows reconstructions produced by the proposed nonlinear residual MLP decoder. Across subjects and image categories, the nonlinear decoder generally produces reconstructions with improved semantic consistency and object recognizability compared with the linear baseline, although improvements in fine-grained textures and low-level visual details remain limited. Ground-truth images are drawn from the publicly available MS COCO dataset [12].

Importantly, the observed improvements were consistent across all evaluated NSD participants. Although absolute reconstruction quality varied between subjects due to individual differences in fMRI signal quality and cortical organization, nonlinear decoding consistently improved semantic alignment relative to linear regression.

### 4.3 Distributional Alignment

We evaluate distributional alignment using MMD and visualize latent structure via PCA (Fig. 6). For diffusion latents *z*, linear decoding is sufficient to match the ground-truth distribution (0.1213 vs 0.1327), and the small gap between *z*_*linear*_ and *z*_*MLP*_ indicates minimal structural change. This is reflected in the overlapping PCA clusters. For CLIP embeddings *c*, the MLP substantially improves alignment (0.0424 vs 0.3585) and results in a large shift from the linear decoder, which is consistent with the separation observed in PCA visualization (Fig. 6). The separation in PCA space visualization confirms the nonlinear transformation introduced by the MLP. Interestingly, we observe weak correlation between MMD and reconstruction quality, consistent with prior findings that distribution-level similarity does not necessarily reflect perceptual or semantic reconstruction quality [23].

**Fig. 6:**
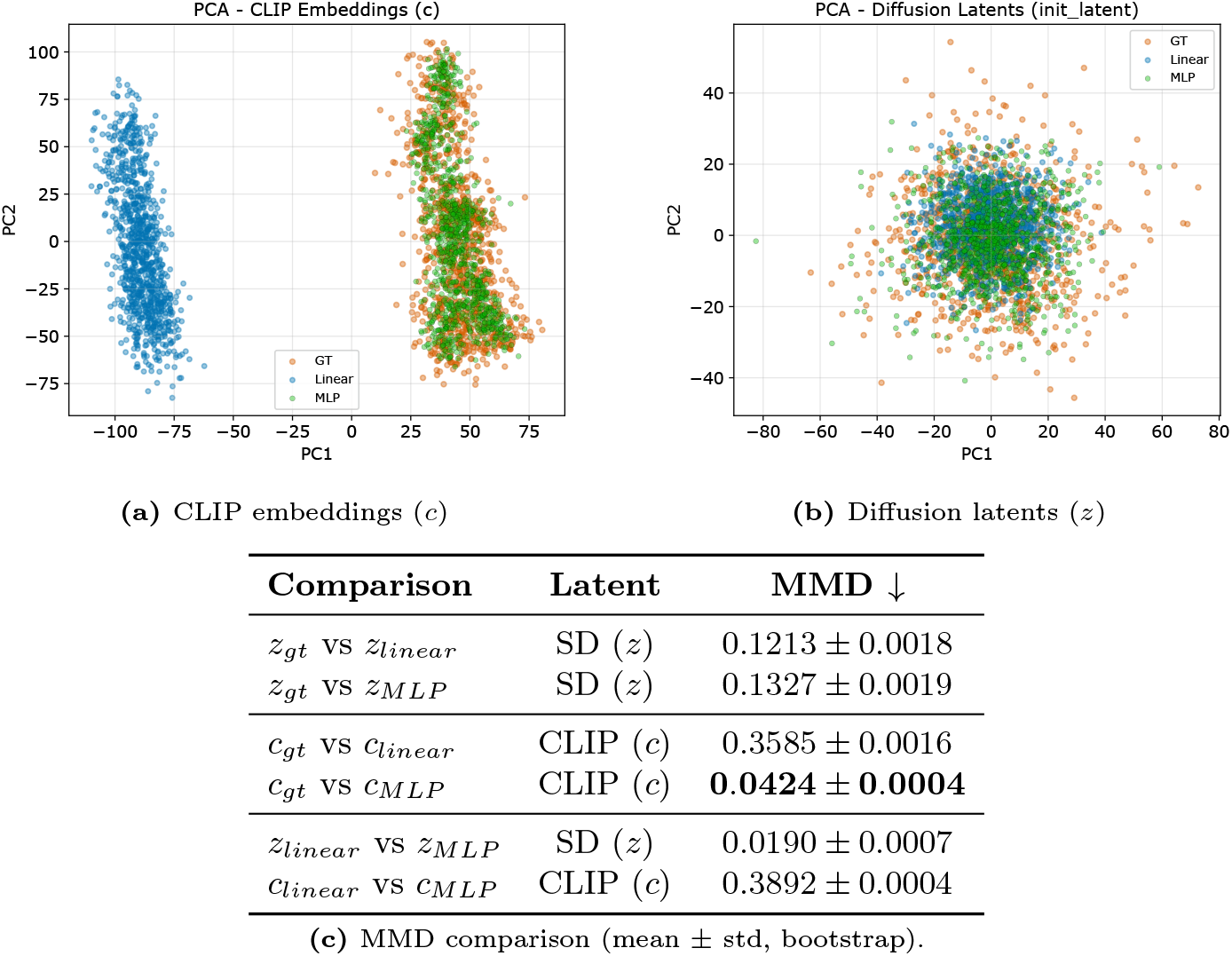
Distribution alignment across latent spaces. (a-b) Top: PCA projections of ground truth (GT), linear, and MLP predictions. (c) Bottom: MMD quantifies distribution mismatch (lower is better). MLP aligns with CLIP embeddings but deviates from diffusion latents, while linear decoding preserves diffusion structure but fails for CLIP. Distributional analyses are shown for subj01 as a representative example.

### 4.4 Cortical Localization of Alignment Gains

We visualize where nonlinear decoding improves brain-to-latent alignment across the cortex. Figure 7 indicates a clear hierarchical structure consistent with our hypothesis: (1) Early visual cortex (V1–V3): improvements are small and spatially sparse, indicating that mappings from voxel activity to diffusion latents are largely linear, consistent with the modest quantitative gains for structural latent prediction. (2) Ventral visual regions (V4, LO, PHC): strong and spatially coherent improvements follow the visual processing pathway, indicating that nonlinear transformations are required to align these representations with semantic embedding spaces. This spatial dissociation provides direct evidence for hierarchical brain-to-latent alignment: early visual areas encode information well captured by linear mappings, whereas higher-order areas encode abstract semantic features requiring nonlinear decoding. These findings also explain the observed discrepancy between latent prediction and reconstruction quality: while nonlinear decoding substantially improves semantic alignment in CLIP space, the resulting distribution shifts limit improvements in pixel-level reconstruction.

**Fig. 7:**
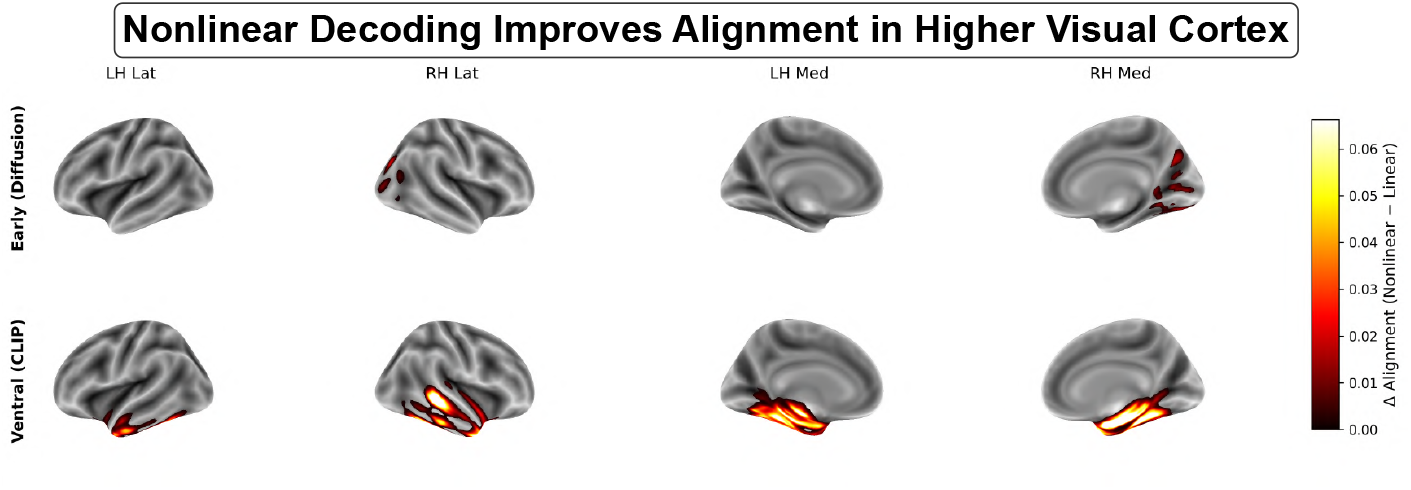
Cortical localization of alignment gains. Change in alignment (*Δ* = non-linear − linear) projected onto cortical surfaces. Top: early visual cortex (V1–V3). Bottom: ventral visual cortex (V4, LO, PHC). Nonlinear decoding yields minimal gains in early areas but strong, spatially coherent improvements along the ventral stream. Representative cortical maps for subj01 are shown

### 4.5 Comparison with Prior Work

As shown in Table 2, Brain2VLM achieves competitive performance across four NSD subjects compared to recent latent diffusion-based methods, including Brain-Diffuser [16], MindLDM [5], and Takagi et al. [20], across both low-level fidelity and high-level semantic metrics. While Brain2VLM achieves a strong performance, these improvements are primarily driven by enhanced semantic alignment rather than substantial gains in pixel-level fidelity. A key distinction lies in the brain-to-latent mapping. Prior works predominantly use linear or ridge regressions to project fMRI activity into latent spaces of pretrained generative models [16, 20], with performance improvements driven by additional conditioning mechanisms rather than improvements in the decoder itself. In contrast, our results show that improving cortical alignment alone leads to consistent gains across metrics (Table 2, Table 4b).

**Table 2.** Quantitative comparison of image reconstruction performance with prior work on the benchmark dataset. Metrics evaluate both low-level fidelity (e.g., PixCorr, SSIM, LPIPS) and high-level semantic alignment (e.g., CLIP, AlexNet, Inception). For feature-based metrics such as CLIP(*l*), AlexNet(*l*), and DINO(*l*), *l* denotes the feature representation extracted from the *l*-th layer of the corresponding pretrained network. ↑ indicates higher is better and ↓ indicates lower is better. Our method, **Brain2VLM**, is compared with BrainDiffuser [16], MindLDM [5], and Takagi et al. [20]

|  | Metric | BrainDiffuser | MindLDM | Takagi et al. | Brain2VLM |
| --- | --- | --- | --- | --- | --- |
| Low-Level | PixCorr ↑ | 0.294 | 0.119 | - | **0.36** |
| | SSIM ↑ | **0.406** | 0.328 | - | 0.31 |
| | LPIPS ↓ | - | - | - | **0.69** |
| | AlexNet(5) ↑ | 87% | **92%** | 83% | **92%** |
| | CLIP(6) ↑ | - | - | - | **71%** |
| | DINOv2(6) ↑ | - | - | - | **69%** |
| High-Level | AlexNet ↑ | - | - | - | **92%** |
| | CLIP ↑ | 63% | 83% | 77% | **85%** |
| | DINO ↑ | - | - | - | **92%** |
| | Inception ↑ | 66% | 81% | 76% | **89%** |
| Retrieval | @1 ↑ | - | - | - | 4% |
| | @10 ↑ | - | - | - | 23% |

Notably, state-of-the-art pipelines employ complex multi-stage architectures while retaining simple regression-based decoding. Brain-Diffuser combines VD-VAE based initialization with dual-guided diffusion using CLIP features [16], while MindLDM integrates MAE-based feature learning, CLIP alignment, VD-VAE depth reconstruction, and ControlNet-based structural guidance [5]. In both cases, only regression models are used in mapping fMRI to latent variables while generative components remain fixed. Similarly, Takagi et al. rely on linear mappings from fMRI to diffusion latents and conditioning embeddings [20].

Despite these architectural complexities, the brain-to-latent interface remains simple. Our results indicate that improving this interface through nonlinear decoding enhances latent prediction quality, which can propagate through the generative pipeline and improve reconstructions.

## 5 Conclusion

We investigated brain-to-latent alignment across four NSD participants and found consistent evidence that decoder expressivity depends on representational hierarchy. Our results suggest that brain-to-latent alignment depends on representational level: early visual cortex can be effectively mapped to structural diffusion latents using linear models, whereas higher-order visual areas require nonlinear transformations to align with semantic embedding spaces, supporting the hierarchical alignment hypothesis. This is consistent with prior work highlighting the nonlinear nature of neural representations and the limitations of simple linear mappings [2,4,22]. The consistency of these findings across subjects suggests that the observed alignment patterns are not specific to an individual participant. We further identify a limitation in current brain decoding systems: improvements in latent prediction, particularly for semantic embeddings, do not reliably translate into improved perceptual reconstruction quality. This highlights a disconnect between representation learning and generative reconstruction in brain-to-image pipelines. These findings suggest that the effectiveness of decoding models depends on the structure of the target latent space, and that improving reconstruction requires balancing latent accuracy with distributional compatibility to the generative prior. Moreover, they provide empirical support for a hierarchical alignment between cortical representations and model latent spaces, consistent with prior work on semantic organization in the visual cortex [7, 10] and recent advances in diffusion-based brain decoding [18, 20].

## 6 Limitations and Future Work

This domain of research can be strengthened in several areas. First, Although the proposed framework generalizes across four NSD participants, larger-scale evaluation on additional datasets and cross-subject decoding remains an important direction for future work. Second, the current decoding framework does not explicitly model latent distributions, and, as observed, can introduce distribution shifts that degrade reconstruction fidelity despite improved latent accuracy. Explicitly modeling these distributions during decoding (e.g., via distribution-aware losses or diffusion priors) may improve compatibility and mitigate the observed trade-offs. Third, the use of relatively simple decoders may restrict the ability to capture complex nonlinear structure in semantic representations, and exploring richer decoders (e.g., transformer-based or multimodal architectures) could better capture these representations. Finally, integrating brain decoding with generative modeling, such as jointly optimizing latent alignment and diffusion dynamics remains an open direction for improving both fidelity and interpretability.

